# Bolstering Wheat’s Immunity: BABA-Mediated Defense Priming Against *Bipolaris sorokiniana* Amid Competition

**DOI:** 10.1101/2024.05.29.596559

**Authors:** Menka Tiwari, Prashant Singh

## Abstract

Plants encounter numerous biotic and abiotic challenges, with biotic stresses significantly limiting wheat productivity. Competition for nutrients and space among plants adds another layer of stress. Defense priming is a promising approach to enhancing plant protection against these environmental stresses. This study explores BABA (β-aminobutyric acid) priming in wheat against *Bipolaris sorokiniana* under varying degrees of competition. We assessed growth parameters, disease phenotype, biochemical changes, and yield-related traits in both primed and non-primed wheat under disease pressure and competition. Our findings revealed that growth parameters declined in both primed and non-primed wheat as competition increased. However, primed wheat showed better morphological growth than non-primed wheat at each competition level. Under disease pressure, primed wheat demonstrated protection comparable to non-challenged plants at all competition levels, while non-primed plants were susceptible. Non-primed wheat under high-density (HD) conditions exhibited the highest disease susceptibility due to intense competition. BABA-primed plants showed better disease protection at each competition level compared to non-primed plants. BABA priming allowed plants to mitigate competition effects and maintain a consistent defense response. The yield performance of primed wheat was superior to that of non-primed wheat across all competition levels. Our research suggests BABA priming as an effective pesticide-free strategy for crop protection against pathogens under competitive conditions.

## Introduction

Plants are constantly exposed to a wide range of biotic and abiotic stressors, with biotic stresses significantly limiting crop productivity in global agriculture. Pathogens disrupt the physiological and biochemical activities of plants, reducing their growth and productivity. However, plants have developed a robust innate immune system to combat environmental challenges, including physical structures like waxy cuticles, thorns, spines, and defensive metabolites (Hilker and Schmülling, 2019). Enhancing the natural immune system of plants offers a safe strategy for plant protection against environmental stresses. One such approach is defense priming, which involves exposing plants to a small dose of stress to enhance their sensitivity and responsiveness to future attacks (Tiwari and Singh, 2021b; Hönig et al., 2023). Primed plants exhibit a faster and stronger defense response due to their prior experience of stress (Mauch-Mani et al., 2017; Tiwari et al., 2022a). Defense priming allows plants to enter an alert or vigilant state, but the defense mechanism does not activate until the plant is challenged with stress (Kerchev et al., 2020). Priming does not involve continuous expression of defense mechanisms, preventing wasteful allocation of photosynthates for defense in the absence of stress (Devi et al., 2023). This smart strategy saves plant resources and contributes to yield enhancement in hostile environments (Tiwari et al., 2024). The priming memory can be maintained throughout the plant’s life, even after the removal of the priming stimulus, and can also be inherited by the progeny generation (Adss et al., 2021; Martínez-Aguilar et al., 2021). Among various methods for enhancing plant resistance, seed priming stands out as an easy, low-cost, and low-risk method for improving growth and tolerance under adverse environmental conditions.

Wheat (*Triticum aestivum* L.) is one of the most extensively cultivated crops globally, serving as a staple food for billions of people. It is the second-largest consumed crop in the world. However, wheat production is limited by various biotic stresses, particularly pathogens (Al-Sadi, 2021). Spot blotch, caused by the hemibiotrophic fungus *Bipolaris sorokiniana*, is one of the most destructive wheat diseases, leading to significant yield losses in warm and humid regions (Devi et al., 2023). The pathogen can persist in contaminated soil, seed, and plant debris, causing up to 100% yield losses (Alkan et al., 2022). Spot blotch is prevalent in South Asia, including China, Nepal, India, and Pakistan, particularly in India’s North Eastern Plain Zone (Gupta et al., 2022). The symptoms of spot blotch are most commonly present on wheat leaves, sheaths, nodes, and glumes, as well as on spikes under severe conditions. The disease manifests as small, light brown lesions on wheat leaves that enlarge and form large patches within a week of infection (Kumar et al., 2020). The conidia of *B. sorokiniana* are thick-walled, elliptical, and have 5–9 septa (Kumar et al., 2020).

Various chemicals, such as BTH, hexanoic acid, pipecolic acid, and azelaic acid, have been used to prime plants against stresses (Tiwari and Singh, 2021a). Recently, β-aminobutyric acid (BABA), a non-protein amino acid, has emerged as a prominent priming agent (Li et al., 2021a). BABA, an isomer of the naturally occurring γ-aminobutyric acid (GABA), is produced by plants under stress conditions (Thevenet et al., 2017). It is a potent inducer of tolerance against both biotic and abiotic stressors (Catoni et al., 2022; Hönig et al., 2023). BABA can be applied through foliar spray, soil drench, and seed treatment. Studies have shown that BABA primes plants to respond robustly to future stress, reducing disease symptoms in various plants, including common bean, tomato, tobacco, and cucumber (Yasir et al., 2023; Martínez-Aguilar et al., 2016; Ramírez-Carrasco et al., 2017; Ren et al., 2022; Kim et al., 2023). BABA priming in *Arabidopsis* resulted in ABA-dependent augmentation of callose formation at infection sites, causing resistance against *Plectosphaerella cucumerina* (Baccelli and Mauch-Mani, 2016). BABA priming decreased the quantity of malondialdehyde in *Vigna radiata* and rice seedlings while increasing proline accumulation and antioxidant enzyme activity during drought and salt stress (Jisha and Puthur, 2016a; Jisha and Puthur, 2016b). Tomato plants primed with BABA via seed priming as well as soil drenching led to the suppression of the nematode *Meloidogyne javanica* by reducing its reproduction rate (Fatemy et al., 2012).

Plant density is a critical factor in agricultural practices, significantly influencing plant growth, development, and interactions with biotic stressors. Higher plant density increases competition for resources like light, water, and nutrients, leading to reduced growth and vigor, making plants more susceptible to biotic stress (Craine et al., 2013; Hol et al., 2013). For example, high plant density enhances competition in maize, resulting in lower biomass output (Zhai et al., 2018). In wheat, increased plant density intensifies intraspecific competition, resulting in reduced tiller earing rate and dry biomass (Xin et al., 2020). Studies have shown that wheat plants grown at higher densities are more susceptible to foliar pathogens due to limited resources for defense compound production (Newton et al., 2010). Environmental stresses and competition in plants play a vital role in shaping plant communities and species dispersion (Zhang and Tielbörger, 2020). Plants undergo various adjustments in response to competition and other environmental stresses (Wang et al., 2021). Increased plant density reduces the horizontal growth of plant roots due to competition for resources in the upper layer of soil but increases the vertical growth of plant roots to acquire resources at greater depths (Wang et al., 2021).

The interplay between plant density and biotic stress response involves trade-offs between resource competition and collective defense mechanisms. Understanding these dynamics is crucial for improving crop yield and ensuring sustainable agricultural practices. BABA priming has already been studied in various plants against biotic and abiotic stressors. In the current study, we examined BABA priming in wheat facing different levels of competition against *Bipolaris sorokiniana* for the first time. We characterized the morphological traits, disease phenotypes, biochemical responses, and yield parameters of BABA-primed and non- primed wheat under varying competition levels. This article outlines BABA-mediated plant defense priming in the context of interactions between plant density and biotic stress response, aiming to maximize plant health and productivity under competitive conditions.

## Materials and methods

### Materials

The wheat seeds (HUW-510) used in the present study were procured from the Department of Genetics and Plant Breeding, Institute of Agricultural Sciences, Banaras Hindu University (BHU), Varanasi. This variety is susceptible to *Bipolaris sorokiniana*, which causes spot blotch disease. HUW-510 is a late-sown variety that was released in 1999/2000. *B. sorokiniana* strain HD3069 (accession no. MCC 1572) was obtained from the Department of Mycology and Plant Pathology, Institute of Agricultural Sciences, BHU.

## Methods

### Seed priming and plant growth

Uniform-sized wheat seeds were selected and surface sterilized using a 1% sodium hypochlorite solution. The seeds were submerged in the sodium hypochlorite solution for 5 minutes and then rinsed three times with sterilized distilled water. Next, the seeds were immersed in a 1 mM BABA solution for 12 hours, after which they were washed three times with sterilized distilled water and allowed to germinate on moist filter paper. The germinated seeds were sown in plastic pots (18 cm in diameter) containing sterilized soil, with varying plant densities in each pot. Three different levels of plant density were used: low density (LD), with 5 plants per pot; medium density (MD), with 10 plants per pot; and high density (HD), with 20 plants per pot (Supplementary Figure 1). The plant density was gradually increased from low to high to enhance the level of competition. The pots were placed in a growth chamber set to a 25/20°C day/night temperature cycle, with a photoperiod of 16 hours of light and 8 hours of darkness, 70% relative humidity, and a photon flux density of 300 µmol mL² sL¹.

### Estimation of morphological parameters

Plants were carefully uprooted 30 days after sowing to estimate various morphological parameters. The roots were thoroughly washed with water to remove the soil. Shoot length and root length were measured using a centimeter scale. Leaf area was measured using a leaf area meter (SYSTRONICS leaf area meter 211). Shoot dry weight and root dry weight were determined by separately oven-drying the shoot and root at 50°C for 48 hours.

### Culturing of pathogen, *B. sorokiniana*

*Bipolaris sorokiniana* was cultured on potato dextrose agar (PDA) media. A 5 mm culture bit was cut from the mother culture plate and placed in the center of a fresh PDA plate. The inoculated plate was then incubated at 25 ± 2°C for 15 days until spore formation occurred.

### Preparation of conidial suspension of *B. sorokiniana* and artificial inoculation of wheat plant with *B. sorokiniana* after 30 days of sowing

The conidial suspension of *Bipolaris sorokiniana* was prepared from a 15-day-old culture plate. The plate, containing black spores, was filled with sterilized water and gently scraped with a sterilized spatula to release the spores without disturbing the media. The suspension was filtered through a double-layered muslin cloth, and the concentration was adjusted to 3 × 10L spores/ml. Tween-20 was added as a surfactant before spraying the conidial suspension onto the wheat leaves. Both primed and non-primed wheat plants, grown at different plant densities (LD, MD, and HD), were then challenged with the pathogen. To maintain humidity, the pots were covered with transparent plastic after inoculation. Symptoms were monitored and recorded after 7 days, and leaves showing symptoms were examined under a compound microscope.

### Disease assessment

The standard area diagram (SAD) was prepared using infected leaves showing various levels of symptoms No symptom – Level 0, 1 to 20% - Level 1, 20 to 40% - Level 2, 40% to 60% - Level 3, 60% to 80% - Level 4, 80% to 100% - Level 5. The severity of the disease was assessed using the SAD, and the percent disease index (PDI) was calculated using the formulae provided by Tiwari and Singh (2024).

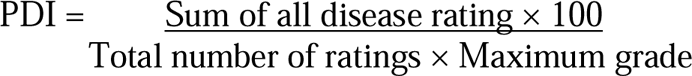

### Biochemical parameters

Leaves from both primed and non-primed wheat plants were sampled seven days after inoculation. Various biochemical parameters, including chlorophyll, carotenoid, malondialdehyde, total phenol, ascorbic acid, and total ROS scavenging activity, were quantified following methods described in our previous research (Tiwari et al., 2022b). Chlorophyll levels were determined using the method outlined by Maclachlan and Zalik (1963), while carotenoids were measured following the procedure by Duxbury and Yentsch (1956). Lipid peroxidation, indicative of membrane damage, was assessed by quantifying malondialdehyde (MDA) content, following the method of Heath and Packer (1968). Total phenolic content was estimated using the method described by Bray and Thorpe (1954), and ascorbic acid content was determined following the procedure outlined by Keller and Schwager (1977). The total ROS scavenging activity was quantified using the method outlined by Blois (1958).

### Proline

The proline content was determined using the acid-ninhydrin method (Bates et al., 1973). 100 mg of leaf tissue was homogenized in 5 mL of 3% (w/v) sulfosalicylic acid and centrifuged at 10,000 rpm for 20 min to separate the extract. The reaction mixture consisted of 1 mL of supernatant, 1 mL of glacial acetic acid, and 1 mL of 0.2% (w/v) ninhydrin, and was incubated for 1 hour at 100°C. The reaction was halted using an ice bath, and the resulting color was extracted using 2 mL of toluene. The absorbance was measured at 520 nm. Free proline was quantified using L-proline as a standard.

### Superoxide dismutase (SOD)

The SOD enzyme activity was determined following the procedure outlined by Fridovich (1974). 0.2 g of leaf tissue was crushed in 2 ml of 100 mM sodium phosphate buffer (pH 7.5) containing 0.5 mM EDTA. The enzyme extract was separated from leaf debris by centrifugation at 10,000 rpm for 20 minutes. A reaction mixture (3 ml) containing 100 mM sodium phosphate buffer (pH 7.8), 2.25 mM nitro blue tetrazolium chloride, 3 mM EDTA, 200 mM methionine, 60 µM riboflavin, and 0.1 ml of the enzyme extract was prepared in a test tube. The reaction mixture was placed at 25°C and exposed to a 400 W bulb for 30 minutes. The absorbance was measured at 560 nm, and the SOD activity was expressed as units per gram of fresh weight (unit g^−1^ fresh wt.)

### Catalase (CAT)

The catalase activity was determined following the procedure outlined by Aebi (1984). For enzyme extraction, 0.2g of fresh wheat leaf was crushed in 5 ml of EDTA phosphate buffer and centrifuged at 10,000 rpm for 20 minutes. The reaction mixture (3 ml) consisted of 0.04 mM H_2_O_2_ in 100 mM phosphate buffer (pH 7.0) and 100 µl of enzyme extract. Catalase (CAT) enzyme activity was assessed by measuring the decrease in H_2_O_2_ absorbance at 240 nm. The catalase activity was expressed as nanomoles of H_2_O_2_ oxidized per minute per gram of fresh weight (n mole H_2_O_2_ oxidised min^−1^ g^−1^ fresh wt.).

### Ascorbate peroxidase (APX)

The APX enzyme activity was determined following the procedure outlined by Chen and Asada (1989). The crude enzyme was extracted by crushing 0.2 g of fresh leaf in 5 ml of EDTA phosphate buffer, and the homogenate was centrifuged at 10,000 rpm for 20 minutes. A reaction mixture (3 ml) containing 100 mM potassium phosphate buffer (pH 7.0), 5 mM ascorbate, 0.5 mM H_2_O_2_, and 600 µl of enzyme extract was prepared. The decrease in optical density of the reaction mixture was measured at 290 nm. APX activity was calculated as nanomoles of ascorbate oxidized per minute per gram of fresh weight (n mole ascorbate oxidised min^−1^ g^−1^ fresh wt.).

### Defence enzymes

The activity of the defense enzymes phenylalanine ammonia lyase (PAL) and peroxidase (POX) was quantified following the methods described in our previous research paper (Tiwari et al., 2024). PAL activity was measured using the method outlined by Dickerson et al. (1984), while POX activity was determined following the procedures outlined by Hartee (1955) and Hammerschmidt (1982).

### Determination of yield-related traits

Yield-related traits of both primed and non-primed wheat were measured after harvesting the spikes. Various parameters including the total number of spikes, spike length, spikelet density, spikelet fertility, grain length, grain width, and thousand grain weight (TGW) were determined following the methods outlined in Tiwari et al., 2022b.

### Statistical analysis

All experiments were performed using three independent biological replicates. The significant impact of BABA-mediated priming on morphological parameters, PDI, biochemical parameters, and yield-related traits was identified using one-way analysis of variance (ANOVA) with the scientific software SPSS 21.0 (IBM Corp, New York). Tukey’s post hoc test was subsequently employed to determine the levels of significant differences among primed and non-primed wheat plants under different levels of competition, with a significance threshold set at p < 0.05.

## Results

### Effect of BABA priming on morphological parameters

The shoot length and root length decreased with increasing plant density, indicating higher competition, in both primed and non-primed wheat plants. However, overall, primed wheat exhibited better growth compared to non-primed wheat at each level of competition (Figures 1A and B). Specifically, at low density (LD), the shoot length increased by 9.64% in primed plants compared to non-primed plants. At medium density (MD) and high density (HD), the shoot length was enhanced by 16.04% and 24.71%, respectively, in primed wheat compared to non-primed wheat (Figure 1C). Similarly, root length was also reduced in non-primed plants compared to primed plants with increasing competition level, but primed plants consistently displayed longer roots compared to non-primed plants at all levels of competition. At LD, primed plants had 58.42% longer roots than non-primed plants. At MD and HD, primed wheat root length increased by 36.04% and 42.85%, respectively, compared to non-primed wheat (Figure 1D). Additionally, we assessed total leaf area and found a similar trend. Total leaf area decreased with increasing competition level in both primed and non-primed wheat. However, primed wheat showed better leaf area compared to non-primed wheat at each level of competition. Specifically, at LD, MD, and HD, primed wheat exhibited a 90.95%, 56.87%, and 28.30% increase in leaf area, respectively, compared to non-primed wheat (Figure 1E). Above- and below-ground biomass was also reduced in both primed and non-primed wheat with increasing competition level, but overall, biomass was higher in primed wheat compared to non-primed wheat. Primed wheat showed enhancement in shoot dry weight by 38.88%, 48.57%, and 120% at LD, MD, and HD, respectively, compared to non-primed wheat (Figure 1F). Similarly, primed wheat exhibited an increase in root dry weight by 70.58%, 53.57%, and 30.95% at LD, MD, and HD, respectively, compared to non-primed wheat (Figure 1G).

**Figure 1.**
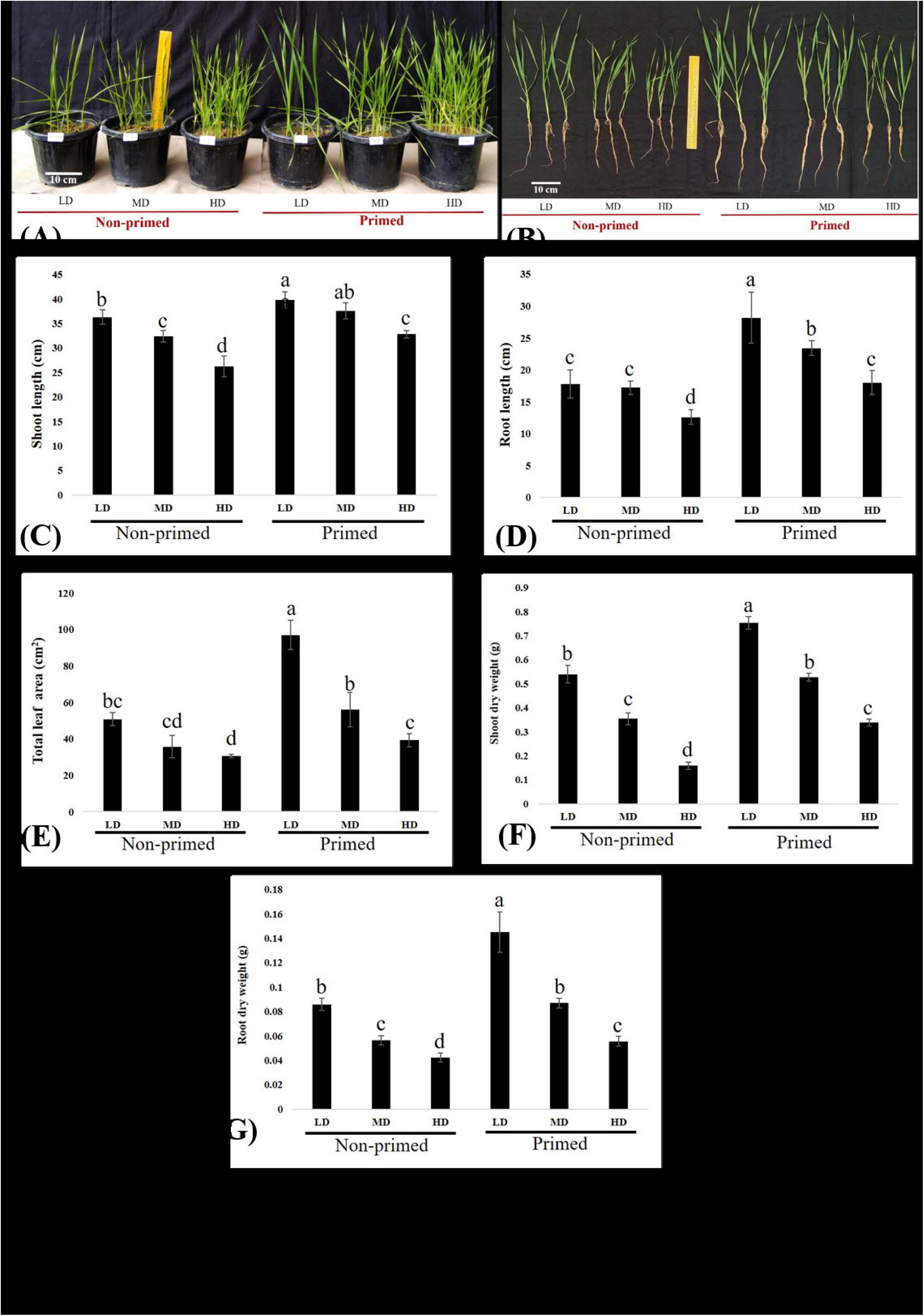
Growth parameters of BABA-primed and non-primed wheat at different plant densities (LD = low density, MD = medium density, HD = high density). **(A)** Plant height, **(B, C)** Shoot length, **(B, D)** Root length, **(E)** Total leaf area, **(F)** Shoot dry weight, **(G)** Root dry weight. Data in C, D, E, F, and G are represented as the mean ± SD. Different alphabets indicate a statistically significant difference at P < 0.05 in Tukey’s test. The results are representative of three independent biological replicates. (n = 20).

### Assessment of disease phenotype in primed and non-primed wheat under varying levels of competition

Assessment of disease phenotypes indicated that primed plants exhibited fewer disease symptoms compared to non-primed plants at each level of competition (Figures 2A and C). The percent disease index was determined using the Standard Area Diagram (SAD) (Figure 2B). Non-primed plants displayed disease index of 76%, 75%, and 93% at low density (LD), medium density (MD), and high density (HD), respectively, while primed plants maintained a consistent disease index of approximately 23% at all competition levels (Figure 2D). The highest disease index was observed in non-primed HD plants (93%). Examination of wheat leaves with symptoms under the microscope confirmed the presence of *Bipolaris sorokiniana* conidia (Figures 2E and F).

**Figure 2.**
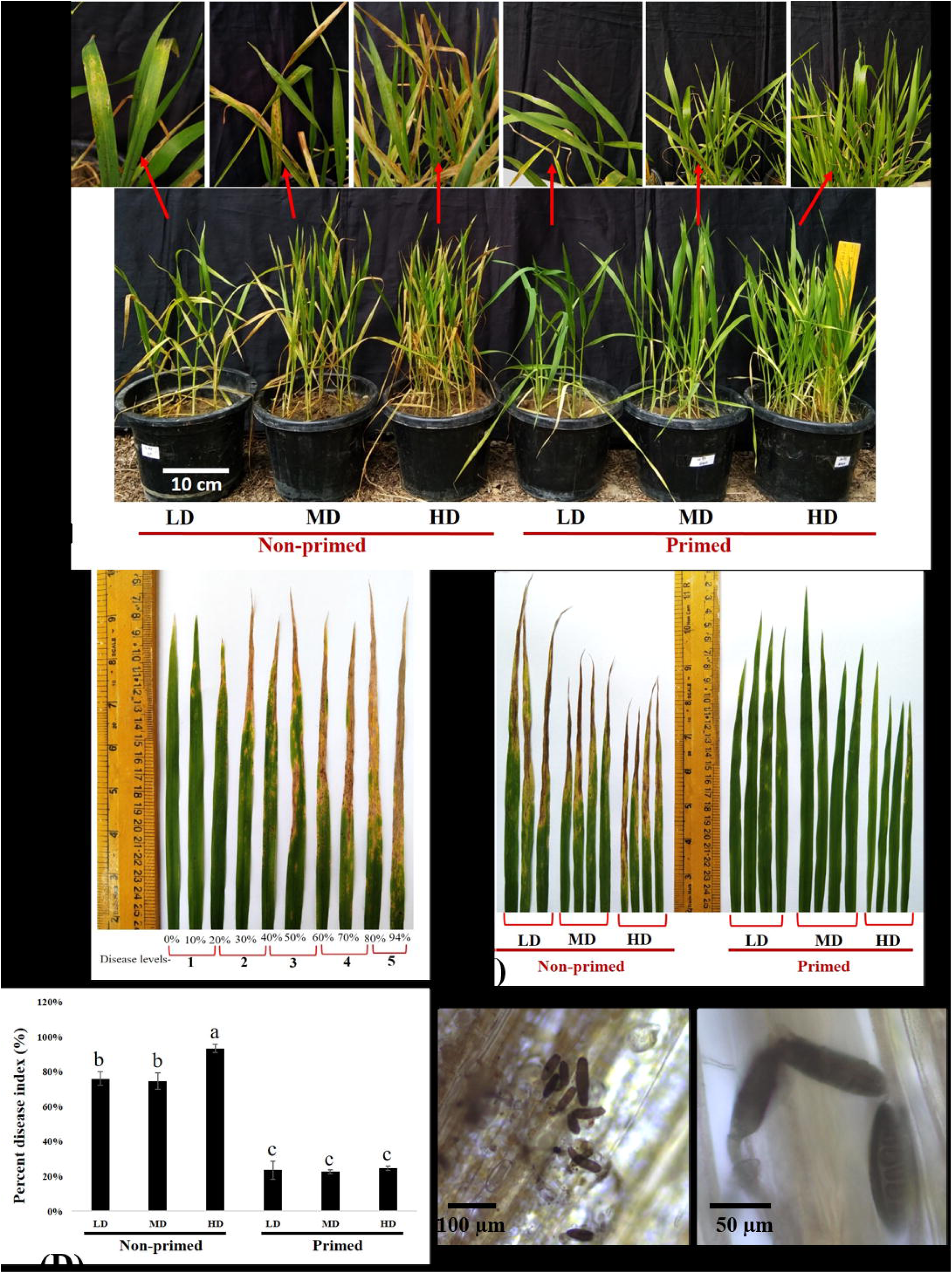
Disease phenotype of spot blotch disease in BABA primed and non-primed wheat at different plant densities (LD = low density, MD = medium density, HD = high density). **(A, C)** Symptoms of spot blotch disease on the leaves of BABA primed and non-primed wheat. **(B)** SAD showing different disease levels. **(D)** Percent disease index of spot blotch in BABA primed and non-primed wheat. **(E, F)** Spores of *Bipolaris sorokiniana* HD3069 on wheat leaves under microscope. Data in D are represented as mean ± SD. Different alphabets indicate statistically significant difference at P < 0.05 in Tukey’s test. Results are representatives of three independent biological replicates. (n = 20).

### Biochemical parameters of primed and non-primed wheat at different levels of competition

Biochemical changes were examined in BABA-primed and non-primed wheat under disease pressure at different levels of competition (Figures 3 and 4). The levels of all metabolites were nearly identical in both BABA-primed and non-primed plants at all competition levels without disease pressure, indicating no direct induction of the defense response due to BABA priming. In non-primed plants under disease pressure, the levels of chlorophyll a and b decreased similarly in both low density (LD) and medium density (MD), with a more pronounced drop in high density (HD). However, when comparing primed and non-primed plants under disease pressure, the levels of chlorophyll a and b in primed LD were elevated by 3.01 and 2.65 times, respectively, compared to non-primed LD (Figure 3A). In primed MD, the levels increased by 2.87 and 3.33 times compared to non-primed MD. In primed HD, the levels of chlorophyll a and b were also enhanced by 8.88 and 5.4 times, respectively, compared to non-primed HD under biotic stress. Moreover, primed plants exhibited similar levels of chlorophyll a and b at all competition levels under disease pressure, which were almost comparable to the non-challenged plants. Similar patterns were observed for other metabolites as well.

**Figure 3.**
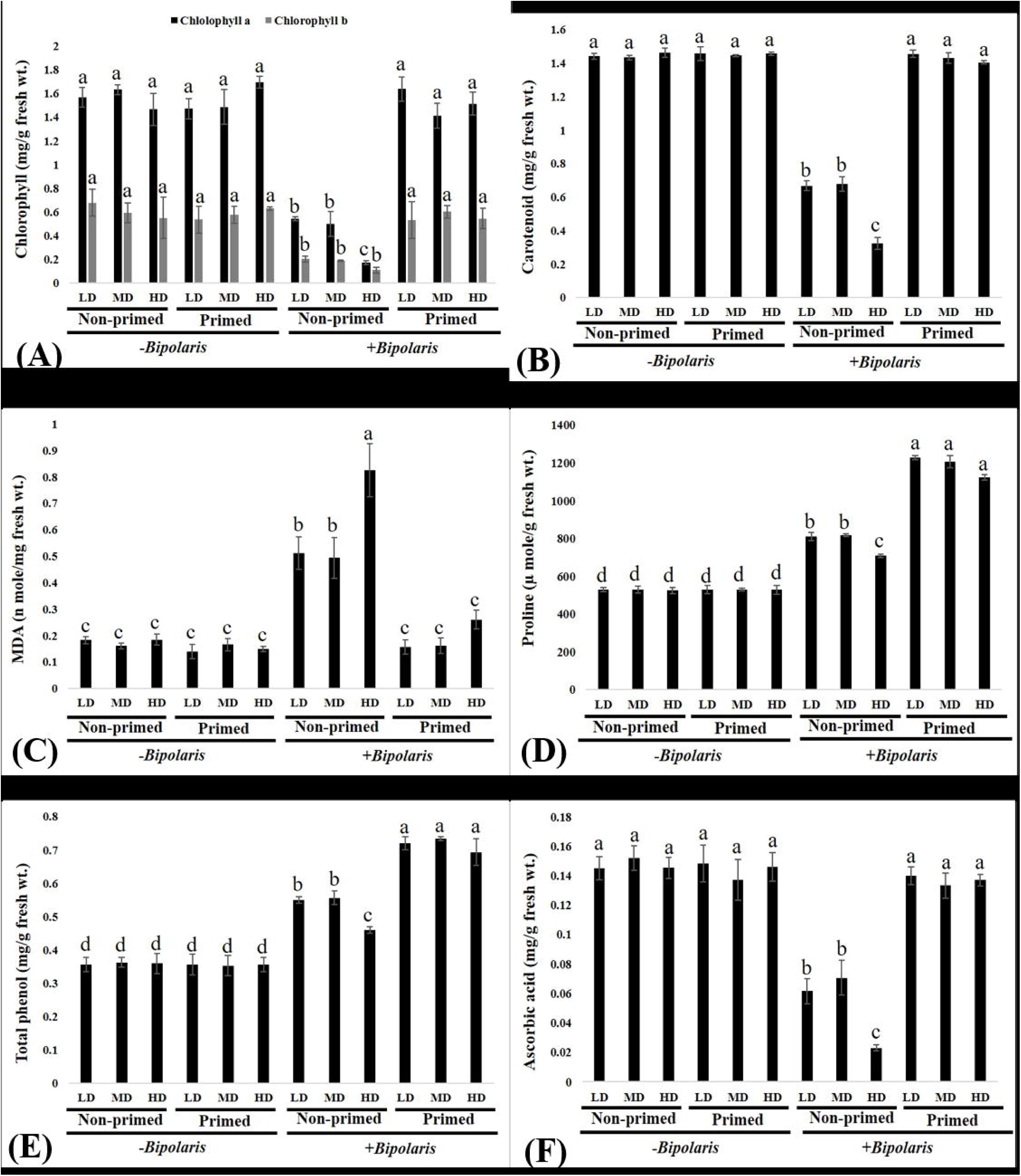
Biochemical parameters of BABA primed and non-primed wheat at different plant densities (LD = low density, MD = medium density, HD = high density) after challenging with *Bipolaris sorokiniana.* **(A)** Chlorophyll a and b, **(B)** Carotenoid, **(C)** MDA content, **(D)** Proline, **(E)** Total phenol and **(F)** Ascorbic acid of BABA primed and non-primed wheat after challenging with *Bipolaris Sorokiniana* HD3069. Data are represented as mean ± SD. Different alphabets indicate statistically significant difference at P < 0.05 in Tukey’s test. Results are representatives of three independent biological replicates. (n = 20).

**Figure 4.**
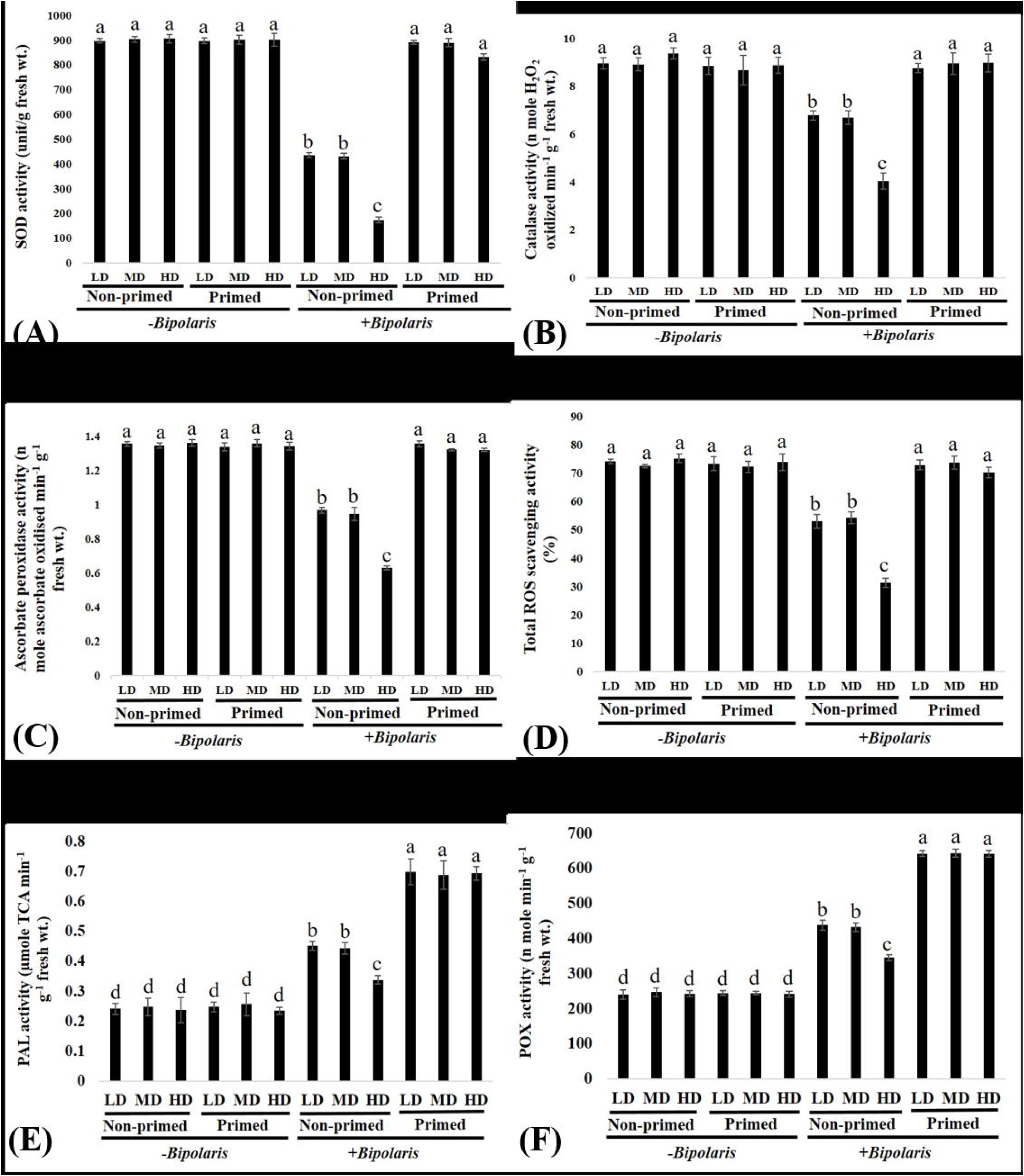
Changes in enzymatic activity and total ROS scavenging activity of BABA primed and non-primed wheat at different plant densities (LD = low density, MD = medium density, HD = High density) after challenging with *Bipolaris sorokiniana.* **(A)** SOD activity, **(B)** Catalase activity, **(C)** Ascorbate peroxidase activity **(D)** Total ROS scavenging activity, **(E)** PAL activity, **(F)** POX activity. Data are represented as mean ± SD. Different alphabets indicate statistically significant difference at P < 0.05 in Tukey’s test. Results are representatives of three independent biological replicates. (n = 20).

The quantity of carotenoid was increased in primed LD, MD, and HD by 2.19, 2.10, and 4.37 times, respectively, compared to non-primed LD, MD, and HD wheat under pathogen pressure (Figure 3B).

We also evaluated the extent of lipid peroxidation in the cell membrane by measuring the MDA content. MDA levels increased in non-primed wheat at all competition levels, with a more pronounced increase in high density (HD) non-primed wheat, indicating greater susceptibility to pathogens. The MDA content was higher in non-primed LD, MD, and HD wheat by 3.24, 3.07, and 3.17 times, respectively, compared to primed LD, MD, and HD wheat (Figure 3C). However, primed plants exhibited almost identical levels of MDA at all competition levels under disease pressure, which were comparable to the non-challenged plants.

Furthermore, we quantified the accumulation of proline and total phenol (non-enzymatic antioxidants) and found that upon pathogen exposure, the levels of both metabolites were enhanced in both primed and non-primed plants at all densities, with a more pronounced increase in primed plants. The amount of proline was increased by 1.51, 1.47, and 1.58 times in primed wheat compared to non-primed wheat at low density (LD), medium density (MD), and high density (HD), respectively (Figure 3D). Additionally, the level of total phenol was also heightened by 0.76, 1.32, and 1.5 times in primed wheat compared to non-primed wheat at LD, MD, and HD, respectively (Figure 3E). Similarly, we assessed the level of ascorbic acid (a non-enzymatic antioxidant) and found that after infection, primed wheat exhibited a 1.51, 1.47, and 1.58 times increase in ascorbic acid compared to non-primed wheat at LD, MD, and HD, respectively (Figure 3F).

Further, we analyzed the enzymatic antioxidants activity of primed and non-primed wheat. SOD activity exhibited a 2.25, 1.87, and 5.95-fold elevation in primed wheat compared to non-primed wheat at LD, MD, and HD, respectively (Figure 4A). Similarly, the activity of CAT demonstrated a 1.29, 1.33, and 2.22-fold increase in primed wheat compared to non- primed wheat at LD, MD, and HD, respectively (Figure 4B). Additionally, APX activity also showed 2.25, 1.87, and 5.95 times augmentation in primed wheat compared to non-primed wheat at LD, MD, and HD, respectively (Figure 4C). Moreover, we assessed the total ROS scavenging activity and found that primed wheat exhibited 1.37, 1.35, and 2.23 times the increase in total ROS scavenging activity compared to non-primed wheat at LD, MD, and HD, respectively (Figure 4D).

The activity of the defense enzyme PAL was enhanced by 1.53, 1.54, and 2.09 times in primed LD, MD, and HD, respectively, compared to non-primed LD, MD, and HD (Figure 4E). Additionally, the activity of POX enzyme was augmented by 1.46, 1.48, and 1.85 times in primed wheat compared to non-primed wheat at LD, MD, and HD, respectively (Figure 4F).

### Yield-related traits of BABA-primed and non-primed wheat under varying levels of competition

We analyzed the yield-related traits of BABA-primed and non-primed wheat under varying levels of competition (Figure 5). The results revealed that, in the absence of disease pressure, primed wheat exhibited enhanced total number of spikes, spike length, spikelet density, and spikelet fertility compared to non-primed wheat at all competition levels. Even when subjected to pathogen challenge, primed plants demonstrated superior yield performance over non-primed wheat at all competition levels. The total number of spikes increased by 1.61, 1.51, and 1.86 times in primed wheat compared to non-primed wheat at LD, MD, and HD, respectively, under biotic stress (Figure 5A). Additionally, spike length was higher by 1.42, 1.40, and 1.63 times in primed wheat compared to non-primed wheat at LD, MD, and HD, respectively, under disease pressure (Figure 5B). Similarly, spikelet density showed an increase of 1.55, 1.55, and 2.24 times, and spikelet fertility was augmented by 1.3, 1.32, and 1.5 times in BABA-primed wheat compared to non-primed wheat at LD, MD, and HD, respectively, under biotic stress (Figure 5C and D).

**Figure 5.**
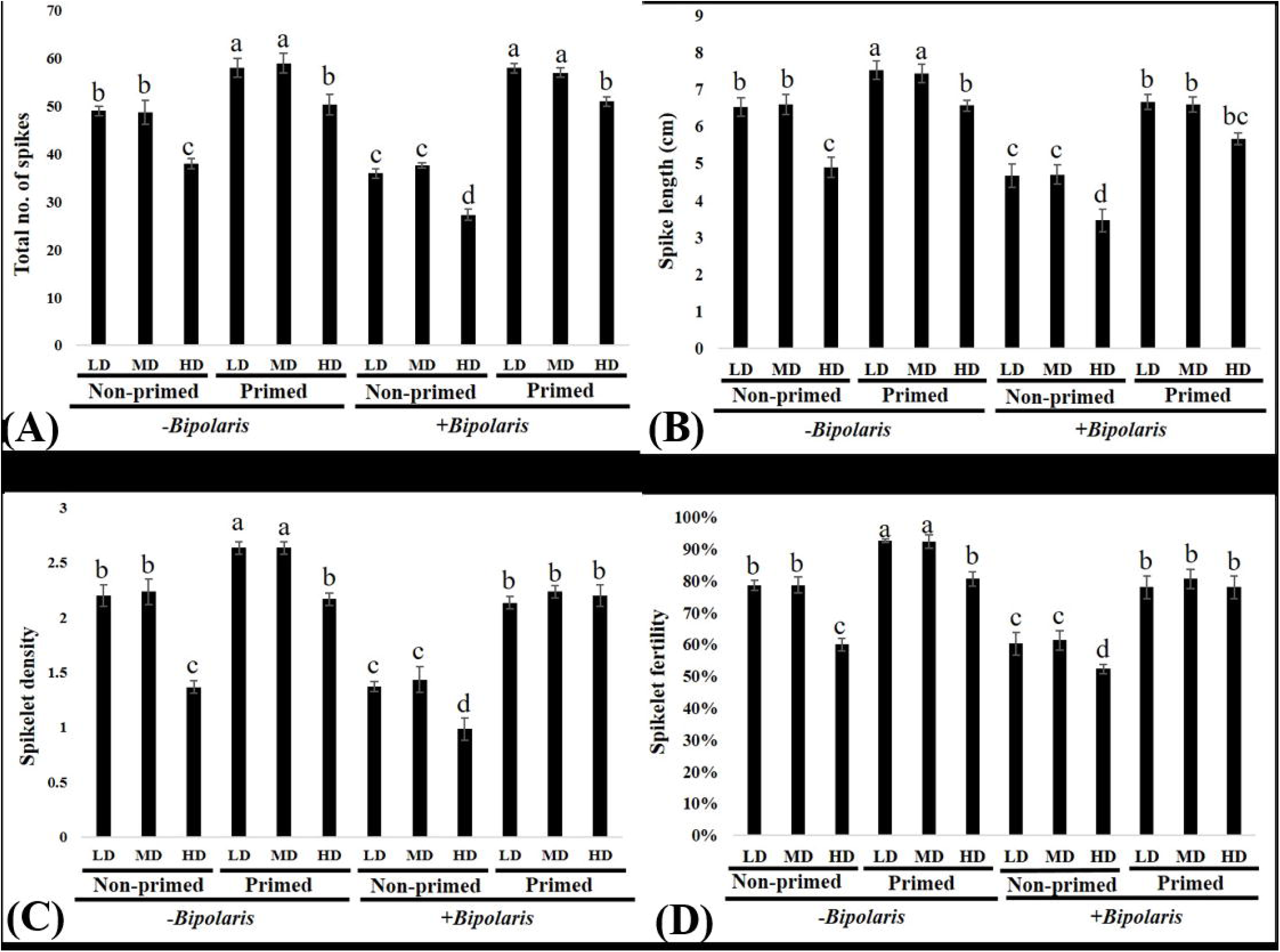
Yield-related traits of BABA primed and non-primed wheat at different plant densities (LD = low density, MD = medium density, HD = High density) after pathogen challenge. **(A)** Total no. of spikes, **(B)** Spike length, **(C)** Spikelet density **(D)** Spikelet fertility. Data are represented as mean ± SD. Different alphabets indicate statistically significant difference at P < 0.05 in Tukey’s test. Results are representatives of three independent biological replicates. (n = 20).

Furthermore, we assessed grain size and weight (Figure 6). The grain width was 1.24, 1.20, and 2.33 times higher in BABA-primed wheat than in non-primed wheat at LD, MD, and HD, respectively, under pathogen pressure (Figures 6A and B). Grain length also increased by 1.09, 1.08, and 1.09 times in primed wheat compared to non-primed wheat at LD, MD, and HD, respectively, after the *B. sorokiniana* challenge (Figures 6C and D). Moreover, the TGW was enhanced by 1.15, 1.17, and 1.35 times in BABA-primed wheat compared to non-primed wheat at LD, MD, and HD, respectively, under disease pressure (Figure 6E).

**Figure 6.**
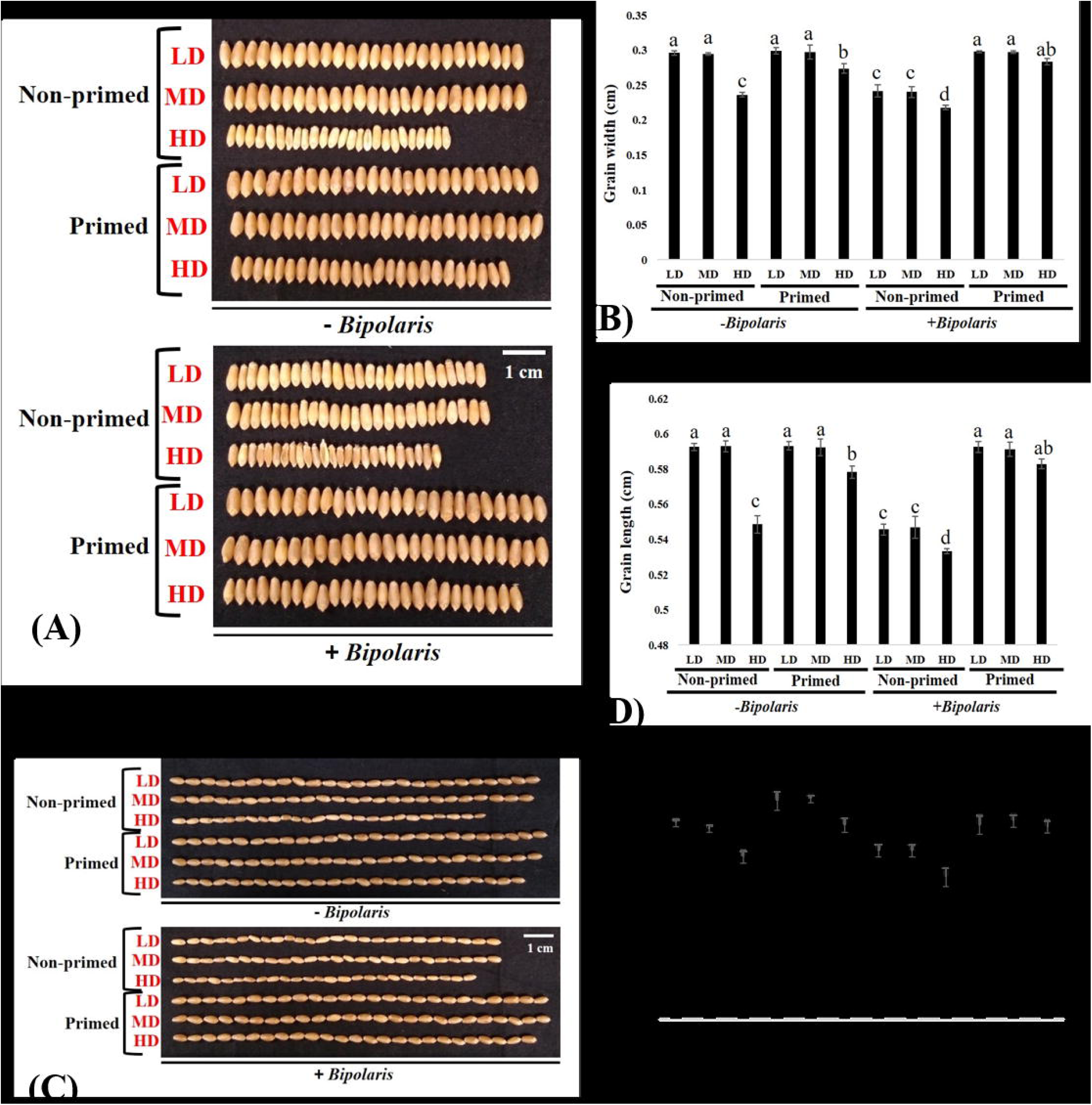
Effect of BABA priming on grain size and grain weight of wheat at different plant densities (LD = low density, MD = medium density, HD = High density) under disease pressure. **(A, B)** Grain width (scale bar = 1 cm, n=25), **(C, D)** Grain length (scale bar = 1 cm, n=25), **(E)** Thousand grain weight. Data in B, D and E are represented as mean ± SD. Different alphabets indicate statistically significant difference at P < 0.05 in Tukey’s test. Results are representatives of three independent biological replicates. (n = 20).

## Discussion

In this study, we observed a decrease in all growth parameters such as shoot length, root length, total leaf area, shoot dry weight, and root dry weight with an increase in the level of competition in both primed and non-primed wheat (Figure 1). This decrease could be attributed to intraspecific competition among wheat plants (Xin et al., 2020). However, BABA-primed wheat exhibited enhanced growth performance compared to non-primed plants at each level of competition. Previous studies have shown that seed priming with BABA enhances shoot length, root length, shoot fresh weight, and root dry weight in chickpeas under salinity stress (Elradi et al., 2022). BABA priming has also been found to promote early seedling growth in bread wheat by enhancing root growth (ÖZKURT and BEKTAŞ, 2022), and to enhance the growth of *Vigna radiata* seedlings under NaCl/polyethylene glycol (PEG) stress (Jisha and Puthur, 2016a). Additionally, BABA priming has been shown to increase the growth of rice seedlings under both stressed and unstressed conditions (Jisha and Puthur, 2016b).

Furthermore, we challenged primed and non-primed plants at different plant densities with *B. sorokininana*. Leaves of non-primed plants exhibited more symptoms and a higher percent disease index (PDI) compared to primed plants at each level of competition (Figure 2). The non-primed plants in HD showed the highest PDI, suggesting that plants facing more competition are more susceptible to the disease. However, primed plants showed almost the same level of PDI at all competition levels, indicating that the effect of competition, initially observed as reduced vegetative growth in primed wheat, is outweighed after biotic stress. The primed plants exhibited a similar level of protection at all competition levels, whether LD, MD, or HD. Exogenous application of BABA has been shown to reduce the level of black shank disease caused by *Phytophthora parasitica* in tobacco (Ren et al., 2022) and significantly restrict the growth of the pathogen *Pseudomonas syringae* pv. tomato DC3000 in *Solanum lycopersicum* (Janotik et al., 2022).

Biochemical changes were assessed in primed and non-primed wheat at various levels of competition. Across all biochemical parameters (including photosynthetic pigments, MDA, proline, phenol, ascorbic acid, SOD activity, catalase activity, APX activity, total ROS scavenging activity, and the activity of defence enzymes PAL and POX), the levels of metabolites were similar in both primed and non-primed wheat at different levels of competition without disease pressure. This suggests that BABA priming does not activate any defense mechanism in the absence of pathogen pressure. Instead, it appears that BABA priming prepares the plants for an enhanced response upon biotic pathogen challenge, rather than directly inducing defense mechanisms.

Examining specific parameters, non-primed wheat exhibited a reduction in chlorophyll and carotenoid levels at all competition levels under disease pressure (Figures 3A and B). The reduction in chlorophyll and carotenoid levels was similar between LD and MD non-primed wheat but more pronounced in HD non-primed wheat, indicating an increased sensitivity to disease with higher competition levels. In contrast, primed plants consistently outperformed non-primed plants under disease pressure at each competition level. Furthermore, within the primed plants under disease pressure, the impact of competition was mitigated, as the levels of chlorophyll and carotenoid were consistent across all competition levels and similar to the non-challenged plants. This suggests that BABA priming enabled plants to counteract the negative effects of competition under disease pressure, as initially observed in their vegetative development. Similar trends were observed in other metabolites, including enzymatic and non-enzymatic antioxidants and defense enzymes. Previous studies have also shown that BABA seed priming can enhance chlorophyll and carotenoid content in rice under Bakanae disease caused by *Fusarium fujikuroi*, possibly due to improved biosynthesis of chlorophyll and reduced breakdown (Gaur et al., 2023; Jisha and Puthur, 2016b).

We also assessed the extent of cell membrane damage caused by *B. sorokiniana* infection by measuring the MDA level. We found that the MDA content was significantly higher in non- primed plants compared to primed plants at each competition level (Figure 3C). However, the increase in MDA level was more pronounced in non-primed HD plants. This indicates that lipid peroxidation of the cell membrane was higher in non-primed wheat than in BABA- primed wheat at each competition level, with the highest impact seen in non-primed HD plants. In contrast, primed plants consistently exhibited similar levels of protection at all competition levels, suggesting that competition did not affect primed plants under disease pressure. Previous studies have shown that BABA-primed tomato plants exhibited reduced MDA content compared to non-primed plants after inoculation with *Alternaria solani*, which causes early blight (Roylawar and Kamble, 2017). Additionally, BABA treatment protected grape berries against postharvest fungus *Botrytis cinerea* by reducing MDA content (Li et al., 2021a).

Furthermore, we quantified the levels of non-enzymatic antioxidants (proline, phenols, and ascorbic acid) in BABA-primed and non-primed wheat at LD, MD, and HD after the *B. sorokiniana* challenge. The levels of proline and phenols increased in both primed and non- primed wheat at each competition level upon pathogen attack, but the increase was higher in primed wheat compared to non-primed wheat at each level (Figure 3D and E). Moreover, primed wheat also exhibited a higher ascorbic acid content compared to non-primed wheat at each competition level (Figure 3F). Notably, among non-primed plants, HD wheat showed the minimum level of non-enzymatic antioxidants (proline, phenol, and ascorbic acid), indicating that HD non-primed plants were under maximum oxidative stress, whereas HD primed plants showed levels of non-enzymatic antioxidants almost comparable to primed LD (proline and phenol) and control (ascorbic acid). This suggests that BABA priming also enables plants to overcome the negative impact of even the highest level of competition.

Proline is an osmoprotectant, and its accumulation helps to maintain the integrity of the cell membrane, osmotic balance, and ROS detoxification against oxidative stress (Aswathi et al., 2023). Exogenous application of BABA increased the production of proline in *Vicia faba* under oxidative stress (Abid et al., 2020). BABA treatment protected peach fruit from Rhizopus rot by elevating the level of total phenolics (Li et al., 2021b). Our current study implies that the accumulation of proline helps to lower the severity of spot blotch disease in BABA-primed wheat plants. Phenols are secondary metabolites that play a defensive role under infection from pathogens (Selim et al., 2022). BABA-induced priming resulted in enhanced production of phenols in mango against anthracnose caused by *Colletotrichum gloeosporioides* (Li et al., 2019). Similarly, the phenol content was also found to be higher in BABA-treated potato plants under the disease pressure of *Phytophthora infestans* (Altamiranda et al., 2008). Ascorbic acid is one of the most abundant antioxidants in plants, which provides the first line of defence against ROS under biotic stress conditions (Boubakri, 2017).

In addition, we measured the activity of enzymatic antioxidants (SOD, catalase, and APX), total ROS scavenging activity, and defence enzymes (PAL and POX) (Figure 4). The activity of all enzymes was similar in both primed and non-primed wheat at all competition levels without disease pressure, indicating the absence of induced resistance due to BABA priming. However, upon pathogen challenge, primed wheat showed enhanced activity of all enzymes compared to non-primed wheat at each competition level. The primed wheat exhibited a consistent level of enzymatic activity at all competition levels under disease pressure, suggesting that competition had no discernible effect on primed plants. The detoxification or scavenging of ROS relies on antioxidant enzymes such as SOD, CAT, and APX. SOD functions by converting superoxide radicals into harmless hydrogen peroxide and molecular oxygen. In contrast, catalase facilitates the conversion of hydrogen peroxide into oxygen and water, while APX is responsible for converting hydrogen peroxide to water (Ighodaro and Akinloye, 2018). Tomato plants treated with BABA showed significantly higher activity of SOD, CAT, and APX when infected with *Alternaria solani* (Roylawar and Kamble, 2017). Wang et al., 2020 reported that BABA priming in grapes led to the upregulation of key genes involved in the enzymatic antioxidant system during *Botrytis cinerea* infection. BABA treatment inhibited Rhizopus rot disease in peach fruit by inducing resistance against the necrotrophic fungus *Rhizopus stolonifera*, which was associated with an increase in PAL enzyme activity (Li et al., 2021b). BABA priming also inhibited green mould disease in orange fruit caused by *Penicillium digitatum* by significantly enhancing PAL and peroxidase enzyme activity. The plants were allowed to set seeds, and yield-related traits were measured.

At maturity, various yield-related traits including total number of spikes, spike length, spikelet density, spikelet fertility, grain width, grain length, and TGW were assessed (Figures 5 and 6). BABA-primed wheat exhibited superior yield performance compared to non-primed wheat at every competition level, both in challenged and unchallenged conditions. Non-primed plants subjected to high competition (HD) showed a significant reduction in these traits. In contrast, primed plants maintained better yield performance even under high competition, with notable improvements in all yield parameters in primed HD. This indicates that BABA priming enabled the plants to withstand both disease and competition pressures simultaneously.

Previous studies have shown that BABA can protect various plant species against a range of environmental stresses and enhance yield parameters (Li et al., 2019). In wheat, BABA treatment combined with other elicitors improved yields by offering protection against *Blumeria graminis* f.sp. hordei and *Rynchosporium commune* (Cohen et al., 2016). Similarly, foliar application of BABA to linseed plants promoted yield-related parameters under stressful conditions (Yasir et al., 2023). The observed enhancement in photosynthetic pigments and enzymatic and non-enzymatic antioxidants, as discussed earlier, likely contributed to the higher yields under pathogen and competition pressures.

Our study highlights the protective effects of BABA priming in wheat against both competition-related stresses and spot blotch disease. This approach demonstrates potential for enhancing crop resilience under biotic stress and ecological challenges, offering a promising tool for crop improvement.

## Conclusions

In our research, we uncovered the remarkable protective effects of BABA treatment on wheat plants, particularly in defending against spot blotch disease even when faced with competition. BABA priming not only enhanced growth parameters under all competition levels but also boosted defense mechanisms against the disease compared to non-primed plants. Intriguingly, primed plants exhibited consistent defense levels across all competition levels, effectively neutralizing the negative impact of competition under biotic stress. Primed plants performed nearly as well as non-challenged plants under disease pressure. Moreover, the benefits of priming extended to yield performance, with primed plants displaying increased yield parameters across all competition levels. Thus, BABA priming equips plants to thrive in competitive environments while effectively combating disease, offering significant advantages even under intense competition.

## Author contributions

PS and MT conceived the idea. MT performed the experiment. MT and PS prepared the manuscript. MT and PS edited and finalised the manuscript. All authors contributed to the article and approved the submitted version.

## Acknowledgments

We would like to thank Prof. Ramesh Chand, Department of Mycology and Plant Pathology, Institute of Agricultural Sciences, B.H.U., Varanasi, for providing us with *B. sorokiniana* HD 3069. We also thank Dr. Sandeep Sharma, Department of Genetics and Plant Breeding, Institute of Agricultural Sciences, Banaras Hindu University (BHU), Varanasi for providing HUW-510 wheat seeds.

---

**Supplementary figure 1.** Wheat seeds were primed with BABA, and both primed and non-primed seeds were sown in the pot with increasing plant density (number of plants per pot). At low density (LD), there were 5 plants per pot; at medium density (MD), there were 10 plants per pot; and at high density (HD), there were 20 plants per pot.

